# Predictive coding of action intentions in dorsal and ventral visual stream is based on visual anticipations, memory-based information and motor preparation

**DOI:** 10.1101/480590

**Authors:** Simona Monaco, Giulia Malfatti, Alessandro Zendron, Elisa Pellencin, Luca Turella

## Abstract

Predictions of upcoming movements are based on several types of neural signals that span the visual, somatosensory, motor and cognitive system. Thus far, pre-movement signals have been investigated while participants viewed the object to be acted upon. Here, we studied the contribution of information other than vision to the classification of preparatory signals for action, even in absence of online visual information. We used functional magnetic resonance imaging (fMRI) and multivoxel pattern analysis (MVPA) to test whether the neural signals evoked by visual, memory-based and somato-motor information can be reliably used to predict upcoming actions in areas of the dorsal and ventral visual stream during the preparatory phase preceding the action, while participants were lying still. Nineteen human participants (nine women) performed one of two actions towards an object with their eyes open or closed. Despite the well-known role of ventral stream areas in visual recognition tasks and the specialization of dorsal stream areas in somato-motor processes, we decoded action intention in areas of both streams based on visual, memory-based and somato-motor signals. Interestingly, we could reliably decode action intention in absence of visual information based on neural activity evoked when visual information was available, and vice-versa. Our results show a similar visual, memory and somato-motor representation of action planning in dorsal and ventral visual stream areas that allows predicting action intention across domains, regardless of the availability of visual information.

## Introduction

Predictions are at the basis of accurate and effective actions. The ability to predict the consequences of a movement allows us to anticipate the future state of our body with respect to the ultimate goal of our action and generate an accurate motor command (Wolpert et al. 1995; Miall and Wolpert 1996). Predictive mechanisms are also crucial to adjust our movements according to rapid changes in the environment and to differentiate the sensory consequences of our own actions from external factors (Blakemore et al. 2000). What information do we use to anticipate actions? Anticipations likely originate from experience. Our everyday-life is rich of actions that elicit visual responses when we see ourselves performing actions, somatosensory responses when we feel our hands moving, and motor responses that generate the action. We can therefore retrieve and use this information to predict how a movement is going to look and feel like.

Recent human neuroimaging work has shown that movement planning affects the activity pattern in areas of the dorsal visual stream (Gallivan et al. 2011; Ariani et al. 2015, 2018) known to be involved in action, as well as the ventral visual stream (Gallivan et al. 2013; Freud et al. 2018), specialized in object perception. However, it still remains poorly understood whether the factors that drive the modulation of the activity pattern in these areas during action planning are related to the visual aspect of the upcoming movement kinematics or to the motor goal signals associated with movement planning. Areas in the dorsal visual stream are likely to have a representation of action planning that can be explained by both motor preparation as well as visual predictions of the impending movement, as the dorsal visual stream is known to be specialized in action (Goodale and Milner 1992). Moreover, neurophysiology studies show that the dorsal stream has strong anatomical connections with somatosensory and premotor areas as well as with the early visual cortex (Luppino et al. 1999, 2003; Passarelli et al. 2011), and processes somatosensory and kinesthetic information associated with passive and active arm movements (Sakata et al. 1973; Mountcastle et al. 1975; Seitz et al. 1991; Lacquaniti et al. 1997; Binkofski et al. 2001; Breveglieri et al. 2002). In addition studies with human participants show that the dorsal stream is involved in visual and tactile exploration of objects (Reed et al. 2005; Dijkerman and de Haan 2007; Konen and Kastner 2008; Marangon et al. 2015). Conversely, the known specialization of areas in the ventral visual stream in perception (Goodale 2014), their strong connections with early visual areas in macaques (Baizer et al. 1991) and humans ((Catani et al. 2003), and their crucial role in the visual identification of shapes (Kourtzi and Kanwisher 2000, 2001; Downing et al. 2001; James et al. 2003), indicate that in these areas, visual anticipations might better explain the representation of action planning as compared to motor preparation.

Although vision for action has been extensively investigated in the past decades (Goodale and Westwood 2004; Grol et al. 2007; Castiello and Begliomini 2008; Rossit et al. 2013; Singhal et al. 2013), much less in known about the neural processing of somatosensory and proprioceptive information during motor preparation and action execution. Yet, these are crucial aspects for skilled motor behavior since they provide us with continuous updates about the state of our movement through online feedback during movement execution. Indeed, movements elicit somatosensory responses that provide us with the awareness of how we move even in absence of visual information. Importantly, the lack of somatosensory and proprioceptive feedback profoundly affects our ability to move. For example, patients who have lost the sense of touch, proprioception, and consequently the awareness of the spatial position of their limbs, have also lost the ability to move spontaneously until they re-learn to do so by constantly monitoring their movements with vision (Cole 1995, 2016; Hermsdörfer et al. 2008). In addition to their crucial role for generating actions, afferent proprioceptive feedback alone modulates motor representations used for cognitive tasks like motor imagery (Mercier et al. 2008).

Kinesthetic memory acquired throughout our life while performing hand actions, enables us to anticipate how a movement is going to feel like even before we initiate it, through somatosensory predictions about the upcoming action. As such, kinesthetic memory likely plays an important role in planning actions that require interacting with our surrounding, such as grasping objects. Indeed, action planning relies on internal models that are based on knowledge of the current position of our hand in space as well as the next state, and the process of getting to the next state involves anticipations of somatosensory consequences of the movement (Miall and Wolpert 1996). To what extent the activity pattern in dorsal and ventral stream areas during action planning is related to anticipating the visual consequences of upcoming actions as opposed to retrieving memory-based information about action performance and motor preparation?

We answered this question using multivoxel pattern analyses on fMRI data to identify the visual, memory-based and motor preparation components for the anticipation of actions. To this end, we manipulated the presence or absence of visual information while participants performed delayed grasp or open-hand movements towards a centrally-located object. While in visual conditions participants relied on continuous visual feedback of the object (during the planning phase) and the hand approaching the object (during the execution phase), in absence of visual information participants relied solely on the memory of the object (during the planning phase) and the somatosensory and motor responses (during the execution phase). Therefore, visual and memory-based anticipations of actions in the planning phase would be reflected in above chance decoding accuracy for the two action types in vision and no vision conditions, respectively. The common aspect between Vision and No Vision conditions were the somatosensory, proprioceptive and kinesthetic components elicited by the two actions, as well as the motor preparation component. In fact, this information differed for the two action types (Grasp vs. Open-hand), but not across visual conditions, as the movements and the object were the same regardless of the availability of visual information. We hypothesize that cortical areas that process the anticipations of somatosensory, proprioceptive and motor components of action would show above chance decoding accuracy for the comparison between action types across visual conditions during the planning phase preceding the action. It seems plausible that areas specialized in perception and visual recognition of shapes, in the ventral visual stream, might have a role in anticipating visual predictions of the upcoming movements. Conversely, areas specialized in action, in the dorsal visual stream, might anticipate the proprioceptive and somatosensory consequences of a movement.

## Materials and Methods

The goal of our study was to investigate the extent to which we could decode action planning in areas of the dorsal and ventral visual stream based on visual, memory-based and somatosensory information. To this aim, in our experiment participants were asked to perform two types of hand actions (Grasp and Open-hand) with the eyes open or closed towards a centrally-located target object (Figure1a). At the beginning of each trial participants were first cued to the action to be performed and whether they had to have their eyes closed or open. Following a delay period, they executed the action (Figure 1b). The delayed paradigm allowed us to analyze the activity patterns during the delay preceding the action and distinguish it from movement execution. We used multi-voxel pattern analysis (MVPA) to examine whether we could decode the dissociation between the two actions (Grasp vs. Open-hand) in areas of the dorsal and ventral visual stream known to be involved in hand actions, such as grasping. In particular, we used independent functional localizers to identify four ventral visual stream areas: the lateral occipital complex (LO), lateral occipital tactile-visual area (LOtv), extrastriate body area (EBA) and the motion sensitive area (MT). In addition, we used an *ad hoc* univariate contrast with the experimental runs to localize five areas in the dorsal visual stream: the anterior intraparietal sulcus (aIPS), ventral premotor cortex (vPM), dorsal premotor cortex (dPM), superior parietal occipital cortex (SPOC) and primary motor/somatosensory cortex (M1/S1).

**Fig. 1.**
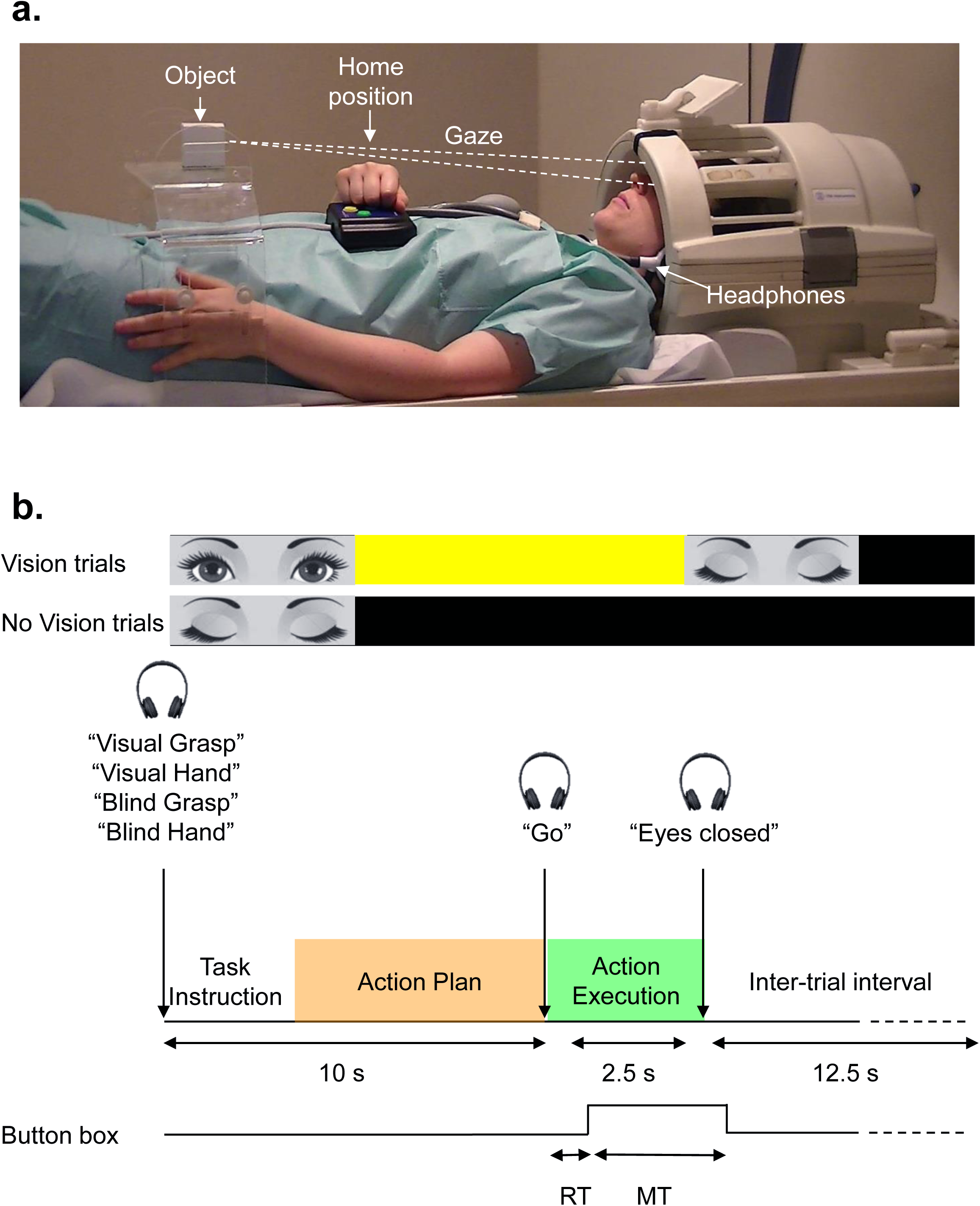
Set-up and experimental paradigm. a) The set-up required participants to maintain direct gaze at a fixation point on the object, when they had their eyes open. The object consisted of a cube and was placed on a platform placed above the participant’s pelvis. After each movement, participants returned the hand on the button box above their abdomen. Participants wore headphone to hear auditory instructions about the task. b) Schematic timing of one trial. At the beginning of each trial, participants heard an auditory cue about the task. The cue was followed by a delay of 10 s, after which a Go cue prompted the participants to perform the action. The next trial started after 12.5 s of intertrial interval (ITI). In Vision trials, participants were instructed to open their eyes at the beginning of the trial until they heard “Eyes closed” at the end of the same trial. In No Vision conditions participants had their eyes closed for the whole duration of the trial. Participants rested their right hand on a button box placed around the navel, and pressed the button until they initiated the movement. We recorded Reaction Times (RTs) and Movements Times (MTs).

### Sessions and subjects

The experimental and localizer sessions took place in two different days. The experimental session lasted approximately 2 hours, including screening and set-up time, while the localizer session took approximately 1.5 hours to be completed.

Nineteen volunteers (age range 23-42, 7 men and 9 women) participated in the experimental session. Sixteen participants also took part in the localizer session. All participants were right handed and had normal or corrected to normal vision.

This study conforms to The Code of Ethics of the World Medical Association (Declaration of Helsinki) printed in the British Medical Journal (18 July 1964). Ethics approval was obtained from the Human Research Ethics Committee of the University of Trento. Informed consent was obtained from all individual participants included in the study.

### Action paradigm

We used high-field (4-Tesla) functional magnetic resonance imaging (fMRI) to measure the blood-oxygenation-level dependent (BOLD) signal (Ogawa et al., 1992) in a slow event-related delayed action paradigm. As shown in Figure 1a, participants performed delayed hand actions using their right dominant hand towards a three-dimensional (3D) target stimulus which was viewed directly without mirrors (Culham et al., 2003). Each trial started with an auditory cue that instructed participants about: i) whether they had to keep the eyes closed or open for the upcoming trial (“Visual” or “Blind”) and ii) the action to be performed at the end of the trial (“Grasp” or “Hand”). The cue was followed by a delay of 10 s after which a Go cue prompted participants to perform the action. The next trial started after 12.5 s of intertrial interval (ITI) (Figure 1b). Grasping movements consisted of reaching-to-grasp the object with a whole hand grip while Open-hand movements required participants to open the hand and move it above the object and touch the object with the palm of the hand but without interacting with the object. Therefore, somatosensory feedback from the object was present in both movement types. In Visual trials, participants were asked to fixate a black dot on the object, therefore the object was in central vision. At the end of each trial, participants returned the hand to the home position, which consisted of a button box placed at a comfortable location around the navel. Participants pressed the button during the ITI by resting their hand on it until they started the next movement. This enabled us to acquire Reaction Times (RT) and Movement Times (MT).

The two by two factorial design, with factors of Action type (Grasp, Open-hand) and Visual Feedback (Vision, No Vision) led to four conditions: Vision Grasp, Vision Open-hand, No Vision Grasp and No Vision Open-hand (Figure 2, upper panel).

**Fig. 2.**
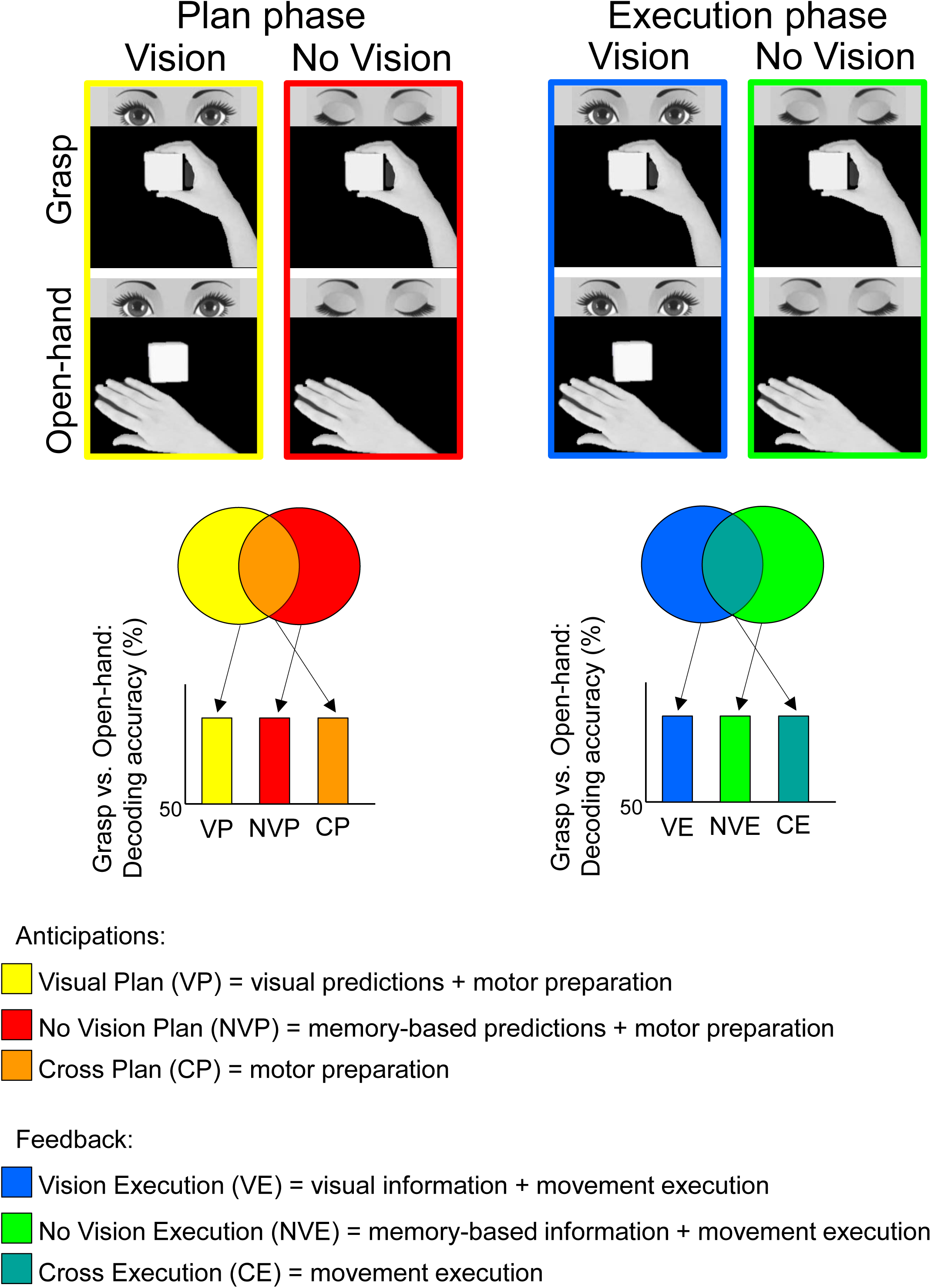
Experimental design. We had a two by two factorial design with two Action types (Grasp and Open-hand) and two Vision conditions (Vision and No Vision), which gave rise to four conditions. We performed our analyses on the Plan (upper left panel) and Execution phase (upper right panel). During the planning phase (lower left panel), the classification between the two action types would allow isolating visual, memory-based predictions and motor preparation. During the execution phase (lower right panel), the classification between the two action types would allow isolating visual and memory based information as well as sensory feedback originating from the execution of the movement.

In our General Linear Model (GLM) we divided each trial in three phases: Task instruction, during which participants heard the auditory cue, Plan phase, during which participants knew what action they were going to perform but were not moving yet, and Execution phase, during which participants performed the action (see *General Linear Model* paragraph for more details). All three phases have been included as predictors in the GLM, however we focused further statistical analyses only on the Plan and Execution phase, as differences in univariate and multivariate analyses within the Task Instruction phase might be explained by information about the auditory instruction per se.

While in Vision trials participants viewed the object throughout the three phases (instruction, plan and execution), in No Vision trials the object was not visible. Visual and memory-based processes, as well as somatosensory anticipations of the upcoming movement would be reflected in above chance decoding accuracy in the planning phase for the dissociation between Grasp and Open-hand movements in Vision, No Vision and across visual conditions, respectively. In particular, while Vision and No Vision conditions might also include a somatosensory component, the cross decoding between visual conditions allowed us to isolate somatosensory anticipations, which are shared in both visual conditions (Figure 2, lower left panel: plan phase). Similarly, visual feedback during the action, memory-based processes as well as somatosensory feedback elicited by the movement itself would be reflected in above chance decoding accuracy for the dissociation between movement types during the execution phase in Vision, No Vision and across visual conditions, respectively. Specifically, while Vision and No Vision conditions include somatosensory feedback during action execution, the cross decoding enabled us to isolate somatosensory feedback and motor commands that elicit responses for both action types regardless of the visual information (Figure 2, lower right panel: action phase).

A video camera placed in the magnet room recorded the eyes of the participants as well as their actions for off-line investigation of the errors. The errors were mistakes in the performance of the task (e.g., eyes open in a No Vision trial, initiating a movement in the plan phase or performing a grasp in an Open-hand condition) and were excluded from further analyses. The 1.8% of total trials were discarded from the analyses because of subject errors.

Each run included 7 trials per experimental condition, for a total of 28 trials per run. Each trial type was presented in counterbalanced order for a run time of ~12,30 min. Participants completed five functional runs for a total of 140 trials per subject (35 trials per condition).

#### Apparatus

Goal-directed actions were performed towards a 3-D stimulus located on a platform secured to the bore bed. The platform was made of Plexiglas and its location could be adjusted to ensure that the participant could comfortably reach the stimulus. The head of the participant was tilted by ~20° to allow direct viewing of the stimuli. The height of the platform could also be adjusted to improve the view and the reachability of the object for each participant. The right upper arm of the participant was supported with foam.

The 3-D stimulus consisted of a white plastic cube (5 cm), which was affixed to the platform with Velcro. The surface of the platform where the object was attached was covered with the complementary side of the Velcro. The platform was placed approximately 10 cm above the subject’s pelvis at a comfortable and natural grasping distance. The stimulus was positioned in the same central location throughout the experiment, therefore the visual presentation of the object remained constant in all trials.

The starting position of the right hand was on a button box placed around the navel while the left hand rested beside the body.

The auditory cues were played through the software Presentation which was triggered by a computer that received a signal from the MRI scanner.

### Localizer paradigms

We ran three sets of localizers to functionally identify ventral stream areas LO, LOtv, EBA and MT. The first two localizers were aimed to identify ventral visual stream areas LO, EBA and MT. Area LO is specialized in object recognition (Kourtzi et al. 2002), while areas EBA and MT are specialized in body-shape recognition and moving stimuli, respectively (Downing et al., 2001; Zeki et al., 1991). For this purpose, participants lay supine in the bed bore and viewed images displayed on a screen through a mirror affixed to the head coil. The third localizer was used to identify LOtv, an object-selective area with visual and tactile properties (Amedi et al., 2001). To this aim, participants tactilely explored objects or textures placed on an apparatus above their torso while fixating a cross viewed through the mirror. While all these areas are specialized in perception, they are also involved in action (Singhal et al., 2011; Monaco et al., 2017; Gallivan et al., 2013). In addition, while LO, EBA and MT are known as visual areas, area LOtv has visual and tactile properties (Amedi et al., 2001). We examined whether the visual and somatosensory specialization of these areas is also present during action planning and execution. We ran *ad-hoc* contrasts to localize these areas based on their specialization (Figure 3a).

**Fig. 3.**
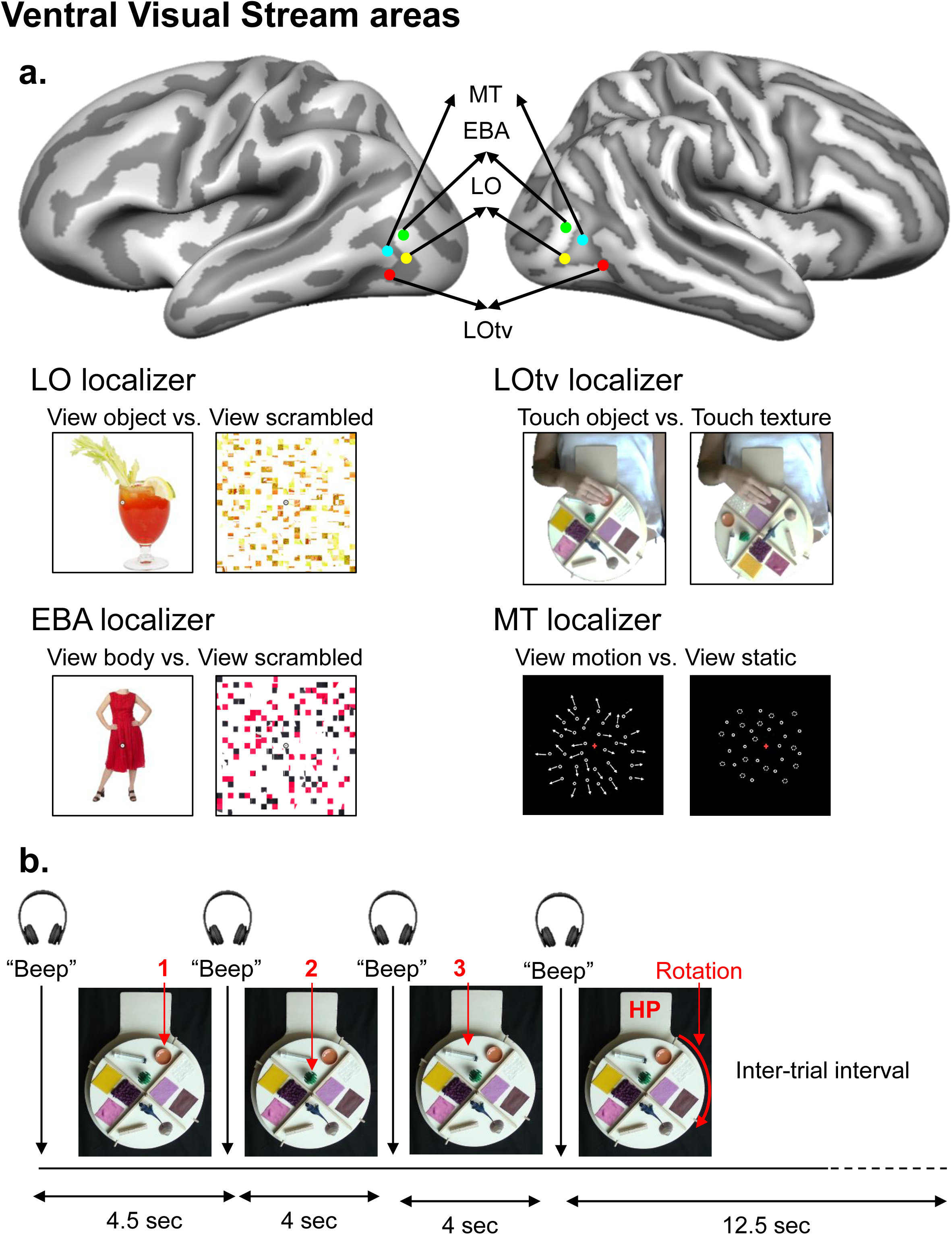
Ventral stream localizers. a) The Talaraich coordinates of area MT, EBA, LO and LOtv are indicated with colored dots. Area LO was localized through a contrast of (Objects > Scrambled images), EBA was identified through a contrast of (Bodies > Scrambled images), MT was localized with the contrast (Motion > Static), and LOtv was identified with the contrast (Touch Object > Touch Textures). b) Apparatus used for the LOtv localizer. Objects and textures were attached on a custom-made wooden turntable and separated into four quadrants by two orthogonal separators. Each quadrant contained three stimuli from one category (objects or textures). In each block of the localizer, participants were required to perform three sequential tactile explorations of the stimuli (objects or textures) in one sector of the turntable. The first beep sound instructed the participant to start to explore the first stimulus on the sector placed next to the right hand (1). Two subsequent beeps separated by 4 s cued participants to start to explore the second and third object (2 and 3). Following 4 s, another beep sound instructed the end of the exploration phase and participants were prompted to return the right hand to the home position (HP) and rotate the wheel with the left hand.

**Fig. 4.**
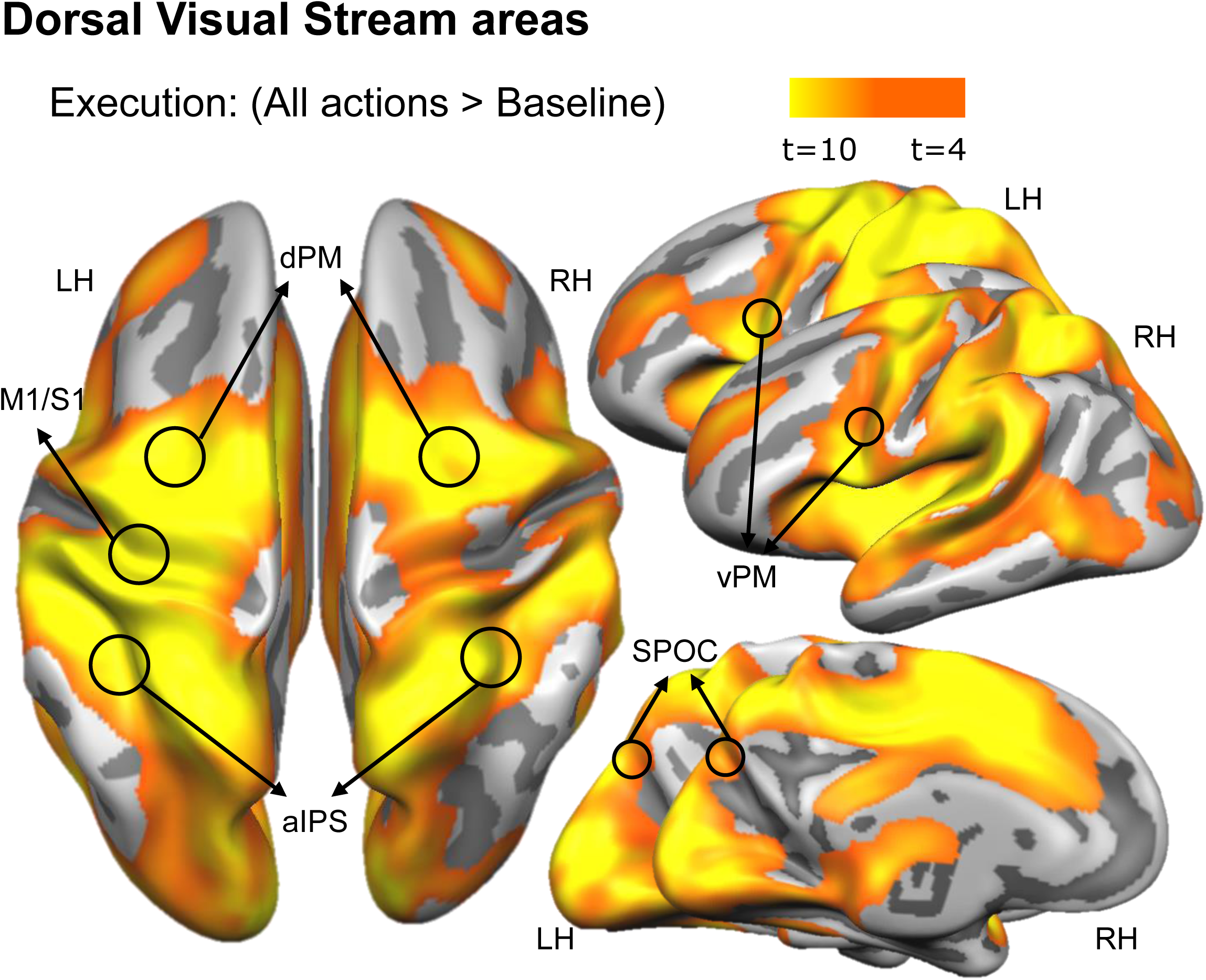
Dorsal stream localizer. Dorsal stream areas were functionally localized by comparing the response during the execution of the two actions to baseline.

#### LO and EBA localizer

The BOT localizer included blocks of color photographs of headless Bodies, Objects and Tools (BOT), as well as scrambled versions of these stimuli (from Gallivan et al., 2013). Stimuli were presented into separate 16-s blocks, with 18 photographs per block, presented at a rate of 400 ms per photograph with a 490-ms inter-stimulus interval. Each run consisted of six stimulus blocks per condition, seven scrambled blocks, and two fixation/baseline blocks (20 s) at the beginning and end of each run. Stimulus blocks were organized into sets of three, separated by scrambled blocks, and balanced for prior-block history within a single run. All subjects participated in two of these localizer runs. Each stimulus block included either three or four repeated photographs, balanced across conditions. Participants were required to fixate a dot that was superimposed on the center of each image and performed an N-back task when the same stimulus was presented twice in a row. The duration of each BOT localizer was 7 min 30 s.

#### MT localizer

Stimuli were presented into 15-blocks of dots moving with center-out trajectories and static dots. Each run included four blocks per condition (Motion and Static) and each Static block was presented after a Motion block and was followed by a 15-s intertrial interval (ITI). The Motion condition consisted of dots moving outwards from the center, while the Static condition consisted of fading non-moving dots. Participants participated in one of this localizer run and were required to fixate a dot placed in the center of the screen for the whole duration of the run. The duration of each MT localizer was 6 min 25 s.

#### LOtv localizer

Participants were required to tactilely explore objects or textures in separate blocks using their right dominant hand. The stimuli were attached on a custom-made wooden turntable placed above the torso of the participant. Two orthogonal separators were mounted on the turntable and divided it into four sectors (Figure 3b). Each sector contained three stimuli from one category (objects or textures). Participants rested the right hand on a flat surface located in middle of the chest and next to the turntable. In each block, participants were required to perform three sequential tactile explorations of the stimuli (objects or textures) in one sector of the turntable for 10.5 s. Blocks were spaced every 12.5 s during which participants rotated the turntable by 45° to bring the next set of stimuli next to the right hand for the subsequent exploration. Therefore, objects and textures were explored in alternating blocks with three items per block. In particular, at the beginning of each block a beep sound instructed participants to start to explore the first stimulus on the sector placed next to the right hand. After 4.5 s, two subsequent beep sounds separated by 4 s, cued participants to start to explore the second and third object. Following 4 s, another beep sound instructed the end of the exploration phase and participants were prompted to return the right hand to the home position and rotate the wheel with the left hand. Stimuli were explored following the same order in all blocks: near-left, far-center and near right stimulus. Participants fixated a fixation cross projected on the screen and viewed through a mirror. Each run included 15 blocks of objects exploration and 15 blocks of texture exploration. Each participant completed three LOtv localizer runs for a total of 45 blocks per condition.

### Imaging parameters

This study was done at the Center for Mind/Brain Sciences (Trento, Italy) using a 4-Tesla 4T Bruker MedSpec BiospinMRscanner and an 8 channel head coil. Functional data was acquired using T2*-weighted segmented gradient echo-planar imaging sequence (repetition time [TR] = 2500 ms; echo time [TE] = 33 ms; flip angle [FA] = 78°; field of view [FOV] = 192 × 192 mm, matrix size = 64 × 64 leading to an in-slice resolution of 3 × 3 mm; slice thickness = 3 mm, 0.45 mm gap). Each volume comprised 35 slices acquired in ascending interleaved order. During each experimental session, a T1- weighted anatomical reference volume was acquired using a MPRAGE sequence (TR = 2700 ms; inversion time TI = 1,020 ms; FA = 7°; FOV = 256×224, 176 slices, 1 mm isotropic resolution).

### Preprocessing

Data were analyzed using the Brain Voyager QX software (Brain Innovation 2.8, Maastricht, The Netherlands). Functional data were superimposed on anatomical brain images, aligned on the anterior commissure–posterior commissure line, and transformed into Talairach space (Talairach and Tournoux 1988). The first four volumes of each fMRI scan were discarded to allow for T1 equilibration. Functional data were preprocessed with spatial smoothing (full-width at half-maximum = 8 mm) and temporal smoothing to remove frequencies below 2 cycles per run. Slice-time correction with a cubic spline interpolation algorithm was also performed. Functional data from each run were screened to ensure that no obvious motion artifacts (e.g., rims of activation) were present in the activation maps from individual participants.

Each functional run was motion corrected using a trilinear/sinc interpolation algorithm, such that each volume was aligned to the volume of the functional scan closest to the anatomical scan. The motion correction parameters of each run were also checked. Data from three participants, two of which also took part in the localizer study, showed abrupt head motion over 1 mm and were discarded from further analyses.

### General Linear Model

In the experimental runs, the group random effects (RFX) GLM included twelve predictors for each participant. There was one predictor for each of the four conditions (Vision Grasp, Vision Open-hand, No Vision Grasp and No Vision Open-hand) by three Phases (Task Instruction, Plan and Execution).. Each predictor was derived from a rectangular wave function (1 volume or 2.5 s for the Task Instruction, 3 volumes or 7.5 s for the Plan phase and 1 volume or 2.5 s for the Execution phase) convolved with a standard hemodynamic response function (HRF; Brain Voyager QX’s default double-gamma HRF). The predictors were: 1) Vision Grasp Instruction, 2) No Vision Grasp Instruction, 3) Vision Open-hand Instruction, 4) No Vision Open-hand Instruction, 5) Vision Grasp Plan, 6) No Vision Grasp Plan, 7) Vision Open-hand Plan, 8) No Vision Open-hand Plan, 9) Vision Grasp Execution, 10) No Vision Grasp Execution, 11) Vision Open-hand Execution, 12) No Vision Open-hand Execution. Although all three phases of the trial (Task Instruction, Plan and Execution) have been included as predictors in the GLM, to answer our questions we focused further statistical analyses on the Plan and Execution phase. We modelled movement parameters (3 rotations and 3 translations), and errors, if present, as predictors of no interest. Contrasts were performed on %- transformed beta weights (β).

### Multivoxel Pattern Analysis

#### Linear Discriminant Analysis (LDA) single-trial classification

MVPA was performed with a combination of in-house software (using MATLAB) and the CoSMo MVPA Toolbox for MATLAB (http://cosmomvpa.org, Oosterhof et al. 2016), with an LDA classifier (http://cosmomvpa.org/matlab/cosmo_classify_lda.html#cosmo-classify-lda). For each participant, we estimated a GLM on non-smoothed data modelling every single trial for each experimental condition. A total of 420 regressors of interest were considered, originating from the four experimental conditions (Vision Grasp, Vision Open-hand, No Vision Grasp and No Vision Open-hand)×three time-phases (Task Instruction, Plan and Execution)×7 repetitions per run×5 runs. In addition, we modelled movement parameters (3 rotations and 3 translations), and errors, if present, as predictors of no interest. We adopted a ‘leave-one-run-out’ cross-validation approach to estimate the accuracy of the LDA classifier. We used beta weights for classification and statistically assessed above chance decoding (50%) significance across participants with a two-tailed one sample t-test.

#### Classifier inputs

To provide inputs for the LDA classifier, the β weights were extracted from the phase of interest (i.e. Plan or Execution phase) for each voxel in the ROI. Each phase included the volumes defined in the predictors for the GLM estimated on unsmoothed data. In particular, the Plan phase consisted of 3 volumes following the Instruction phase, while the Execution phase consisted of 1 volume following the Plan phase.

#### Cross-decoding

We tested whether an LDA classifier trained to discriminate between two conditions could then be used to accurately predict trial identity when tested on a different set of trials (i.e. cross-decode). The trials of the train and test runs were taken from different visual feedback conditions such that the training was performed considering the pairwise comparison between the two actions (Grasp vs. Open-hand) in the No Visual condition and tested in the Visual condition, and vice versa. The cross-decoding accuracies for each subject were computed by averaging together the results obtained in each cross-decoding (i.e., train on No Vision and test on Vision; train on Vision and test on No Vision).

### ROI Analysis

Because we had *a priori* hypotheses about specific areas, we used independent localizer runs to identify regions of interest (ROIs) in the ventral visual stream (LO, LOtv, EBA and MT) and an independent contrast with the experimental runs to localize areas in the dorsal visual stream (SPOC, aIPS, M1/S1, vPM and dPM). The selection criteria based on the localizer runs were independent from the key comparison and prevented any bias towards our predictions (Kriegeskorte et al. 2010; Vul et al. 2010).

We identified each ROI at the group level by using specific contrasts based on the known functional role of each area (see details below). Each ROI consisted of a sphere with radius = 9mm and volume = 3071mm^3^ (or anatomical voxels). We centered the sphere on the voxel with peak activity for the contrast used to localize the area, and in the vicinity of the expected anatomical location of that area. Each sphere included voxels that were selective for the contrast used to define the regions in the dorsal and ventral visual stream. This approach ensured that the same number of voxels was included in each ROI for adequate pattern classification, and that regions were selected objectively and independent of the planned comparisons for pattern classification in the experimental runs. As also shown in previous studies, we found some overlap between ventral stream areas. In particular, we found an overlap between area LO and EBA, MT and EBA (Downing et al. 2001), LO and MT (Kourtzi et al. 2002), LO and LOtv (Amedi et al. 2001) and MT and LOtv. The overlap between LO and EBA was 27% (840 voxels) in the left hemisphere and 6% (200 voxels) in the right hemisphere. Area MT and EBA overlapped by 60% (1832 voxels) in the left hemisphere and 26% (806 voxels) in the right hemisphere. Area LO and MT overlapped by 56% (1718 voxels) in the left hemisphere and 1% (20 voxels) in the right hemisphere. Area LO and LOtv overlapped by 7% (224 voxels) in the left hemisphere, and area MT and LOtv overlapped by 3% (80 voxels) in the left hemisphere.

It is important to note that ROIs that overlap with others are, by definition, non-independent from the regions they overlap with. Despite the fact that our methodological approach, consisting of centering the sphere on the peak-activation voxel, has led to overlaps between adjacent regions, these areas also showed a functional overlap, as previously indicated also in other studies (Downing et al. 2001; Amedi et al. 2002; Kourtzi et al. 2002). The functional overlap between areas makes it difficult to define regions that are completely independent from each other, especially when there is significant anatomical vicinity. There are several reasons why neural systems might overlap as measured with fMRI. First, functionally independent neural populations might occupy the same region of the cortex where neurons that are most selective to function A are intermingled with neurons that are most selective to function B. Second, the dividing line between two functional systems that are spatially segregated but adjacent is unlikely to fall exactly along the voxel borders. Third, the low spatial resolution of fMRI decreases the independence of the signal from bordering voxels. A further contributing factor to the overlap between areas LO and EBA in our study might be related to the fact that both objects and bodies were localized with respect to highly-similar scrambled conditions. Therefore, both ROIs might include non-specific object-form voxels. Another possibility is that there might be a gradient of selectivity, with highest selectivity to objects or body in voxels away from the overlapping zone, and lowest selectivity in voxels within the overlapping zone.

### Localization of ventral stream areas

The visual object-selective area LO was localized by using the contrast (Objects > Scrambled). LO was defined around the peak voxel of activity near the lateral occipital sulcus (Malach et al., 1995; Grill-Spector et al., 1999, 2001). The body-selective area EBA was selected using the contrast (Bodies > Scrambled). The EBA was defined around the peak voxel of activity in the posterior inferior temporal sulcus/middle temporal gyrus (Downing et al., 2001; Peelen & Downing, 2005c), dorsal to LO. The motion-selective area MT was identified with the contrast (Motion > Static) with peak activity in the posterior middle temporal gyrus (Zeki et al. 1991). The tactile-visual object-selective area LOtv was identified with the contrast (Touch Object > Touch Textures) with peak activity located ventrally and anteriorly compared to LO.

### Localization of dorsal stream areas

To localize areas involved in action execution (SPOC, aIPS, M1/S1, vPM and dPM) we use an RFX GLM of the experimental runs to generate an activation map computed by comparing the response during the execution of the two actions to baseline: [(Vision Grasp Action + No Vision Grasp Action + Vision Open-hand Action + No Vision Open-hand Action) > baseline). We selected the peak activation near the expected anatomical location of the ROI: superior end of the parietal occipital sulcus for SPOC (Gallivan et al. 2009; Cavina-Pratesi et al. 2010b; Monaco et al. 2011), junction of the intraparietal sulcus and the inferior segment of the postcentral sulcus for aIPS (Binkofski et al. 1998; Culham et al. 2003; Frey et al. 2005; Begliomini, Caria, et al. 2007; Begliomini, Wall, et al. 2007), omega-shaped area of the central sulcus corresponding to the representation of the hand in M1/S1 (Hari et al. 1993; Yousry et al. 1997), T-junction of the superior precentral sulcus and the caudal end of the superior frontal sulcus for dPM (Cavina-Pratesi et al. 2010; Monaco et al. 2011), inferior part of the junction between the inferior frontal sulcus (IFS) and precentral sulcus for vPM (Tomassini et al. 2007).

### Control areas

To validate the performance of the classifier outside of the ROIs for which we had *a priori* hypotheses, we selected two control areas where activation patterns are not expected to distinguish between actions with nor without vision. To this aim, we reduced the statistical threshold of the activation map resulting from the contrast used for localizing dorsal stream areas (All Actions > baseline) to p = 1 and selected the active voxels within a sphere with radius = 9mm centered on: 1) the white matter adjacent to the right ventricle, and 2) the area just outside of the brain near the left visual cortex (Supplemental Figure 1).

### Information isolated for each condition and phase of the task

#### Cross decoding

The common aspect between Vision and No Vision conditions during the planning phase was the motor preparation for Grasp and Open hand movements. Since proprioceptive feedback is always available, even when we stay still and regardless of visual information, the motor preparation was likely based on the awareness of the current position of the hand relative to the target and the anticipation of somatosensory consequences of the upcoming action. We expected that areas of the dorsal stream, known to be involved in computing movement trajectories and motor planning, would show accurate cross decoding for Grasp and Open hand movements between Vision and No Vision conditions. During the execution of the movement, somatosensory feedback was available in addition to proprioceptive information regardless of the presence or absence of vision.

#### Vision

During the planning phase, vision of the target is available in addition to the proprioceptive information about the current state of our hand. As previously shown, ventral as well as dorsal stream areas show activity patterns for different actions that can be reliably classified in visual conditions (Gallivan et al. 2011, 2013). During the execution phase, participants could rely on somatosensory and proprioceptive feedback as well as vision of the target and the hand approaching the target.

#### No Vision

During the planning phase, participants could rely on memory-information about the target and the proprioceptive information of the current position of the hand. During the execution phase, somatosensory information was also available. Visual-memory processes might also take place to mentally update the estimated position of our hand relative to target, based on somatosensory and proprioceptive feedback. Given the known role of ventral stream areas in memory tasks, and dorsal stream areas in somatosensory and proprioceptive feedback, we expected that both ventral and dorsal stream areas would show activity patterns that could be classified for different actions in No Vision conditions.

### Statistical analyses

#### Behavioural analyses

The RTs were calculated as the time elapsed from the Go cue to button release (Figure 1b, lower panel). The MTs consisted of the duration of a movement measured from button release to button press. It is worth noting that RT and MT coincided with the Action Execution predictor in our GLM. Therefore, behavioural differences between conditions during this phase might be reflected in differences in brain signals detected during the Action Execution phase.

To evaluate differences between conditions, we ran an ANOVA (p<0.05) with repeated measures for each dependent variable (RT and MT) using SPSS. We had a two by two factorial design with two Action types (Grasp and Open-hand) and two Vision conditions (Vision and No Vision), which gave rise to four conditions.

Behavioural data from one participant out of 16 could not be recorded for technical reasons.

#### Multivoxel Pattern analyses

Our initial approach was to use univariate analyses to test our hypotheses on the β weights extracted from each ROI using the group RFX GLM (see Supplemental Materials for details). We then complemented these analyses using a multivariate classification method. The multivariate approach allowed us to isolate the somatosensory anticipation of the planned action as well as the somatosensory feedback during action execution by decoding action type (Grasp vs. Open hand) across visual conditions (train in Vision and test in No Vision condition, and vice versa). In addition, MVPA enabled us to test our hypotheses when activation levels for key conditions did not differ from each other (i.e. Grasp Plan ~ Open-hand Plan) or were at baseline. Indeed, MVPA has the advantage of revealing information carried by the activity pattern in a given area for two (or more) conditions with similar or around baseline activation levels.

For each ROI we performed two-tailed one sample t-tests on the decoding accuracy for the dissociation between Grasp and Open-hand in Vision, No Vision and across visual conditions against chance (50%) during the Plan and Execution phase. To further explore whether the decoding accuracy was higher in Visual conditions, we performed two tailed paired sample t-tests between the decoding accuracy in Vision vs. No Vision, Vison vs. across visual conditions and No Vision vs. across visual conditions.

To control for multiple comparisons, a false discovery rate (FDR) correction of q ≤ 0.05 was applied, based on the number of ROIs and number of t-tests performed within each time phase (Benjamini and Yekutieli 2001).

## Results

### Behavioural analyses

Average reaction and movement times are shown in Supplemental Table 1.

As for RTs, participants were faster to initiate grasp than open hand movements, as well as when vision was available as compared to when it was not. As shown in Supplemental Figure 2, we found a main effect of Action type (F_(1, 14)_ = 8.5, p<0.05) and a main effect of Vision (F_(1, 14)_ = 7.1, p<0.05), but no significant Action type by Vision interaction (F_(1, 14)_ = 2.3, p=0.15).

As for MTs, the durations were comparable across conditions. Specifically, there was no main effect of Action type (F_(1, 14)_ = 3.5, p=0.083), no main effect of Vision (F_(1, 14)_ = 3.1, p=1), nor a significant Action type by Vision interaction (F_(1, 14)_ = 0.3, p=0.6).

### Univariate analyses

As shown in Supplemental Figure 3, the univariate analyses revealed that activations levels were at baseline in most areas during the plan phase, with no interaction between Action and Vision, nor significant differences between Grasp vs Open hand movements, except for four areas that showed a main effect of Action. Therefore, we further explored the patters of activity in our ROIs using the classifier.

### Multivoxel pattern analysis

Figure 5 shows the mean classification accuracy in each ROI for pairwise comparisons of movement type (Grasp vs. Open hand) in Vision and No Vision condition as well as across visual conditions. The Talairach coordinates and numbers of voxels of each ROI are specified in Table 1. Statistical values are reported in Table 2. We report only significant results that survived FDR correction for multiple comparisons.

**Fig. 5.**
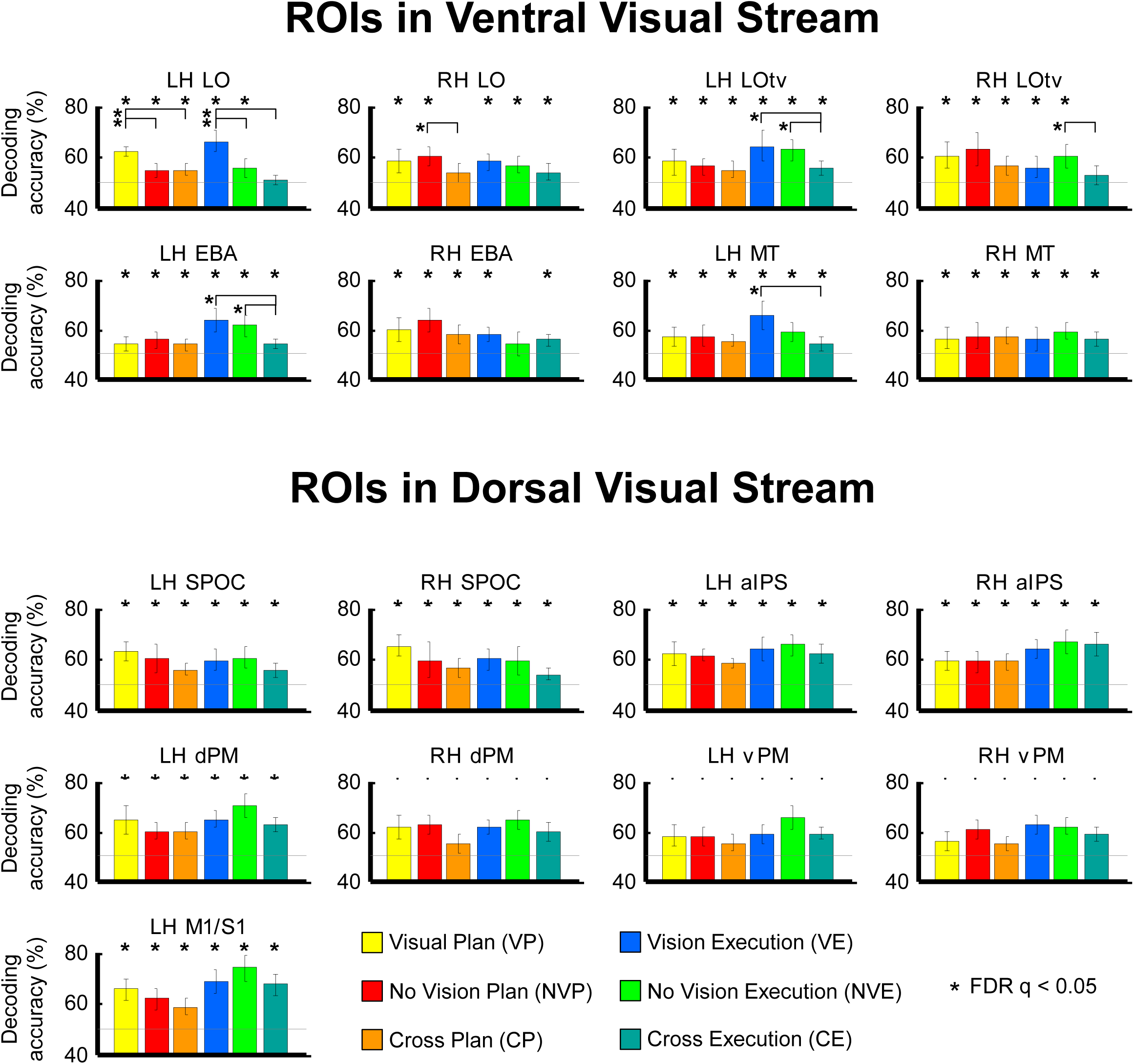
Movement decoding from our ROIs. The plots show decoding accuracies in ventral and dorsal visual stream areas for the dissociation between Grasp and Open-hand movements in Vision and No Vision condition as well as across Visual conditions during the planning phase (yellow, red and orange bars, respectively) and during the execution of the movement (blue, green and turquoise bars, respectively). Chance level is indicated with a line at 50% of decoding accuracy. Error bars show 95% confidence intervals.

**Table 1.** Talairach coordinates for each ROI.

|  | Talairach coordinates |  |  |
| --- | --- | --- | --- |
|  | X | Y | Z |
| LH LO | -37 | -82 | -3 |
| RH LO | 46 | -69 | -7 |
| LH LOtv | -43 | -61 | -11 |
| RH LOtv | 46 | -49 | -9 |
| LH EBA | -48 | -71 | 4 |
| RH EBA | 47 | -70 | 7 |
| LH MT | -44 | -71 | 1 |
| RH MT | 51 | -61 | 7 |
| LH dPM | -30 | -12 | 50 |
| RH dPM | 31 | -12 | 49 |
| LH vPM | -45 | 2 | 14 |
| RH vPM | 42 | -2 | 12 |
| LH M1/S1 | -34 | -32 | 48 |
| LH aIPS | -37 | -43 | 44 |
| RH aIPS | 38 | -39 | 44 |
| LH SPOC | -15 | -73 | 32 |
| RH SPOC | 16 | -73 | 36 |

**Table 4.**
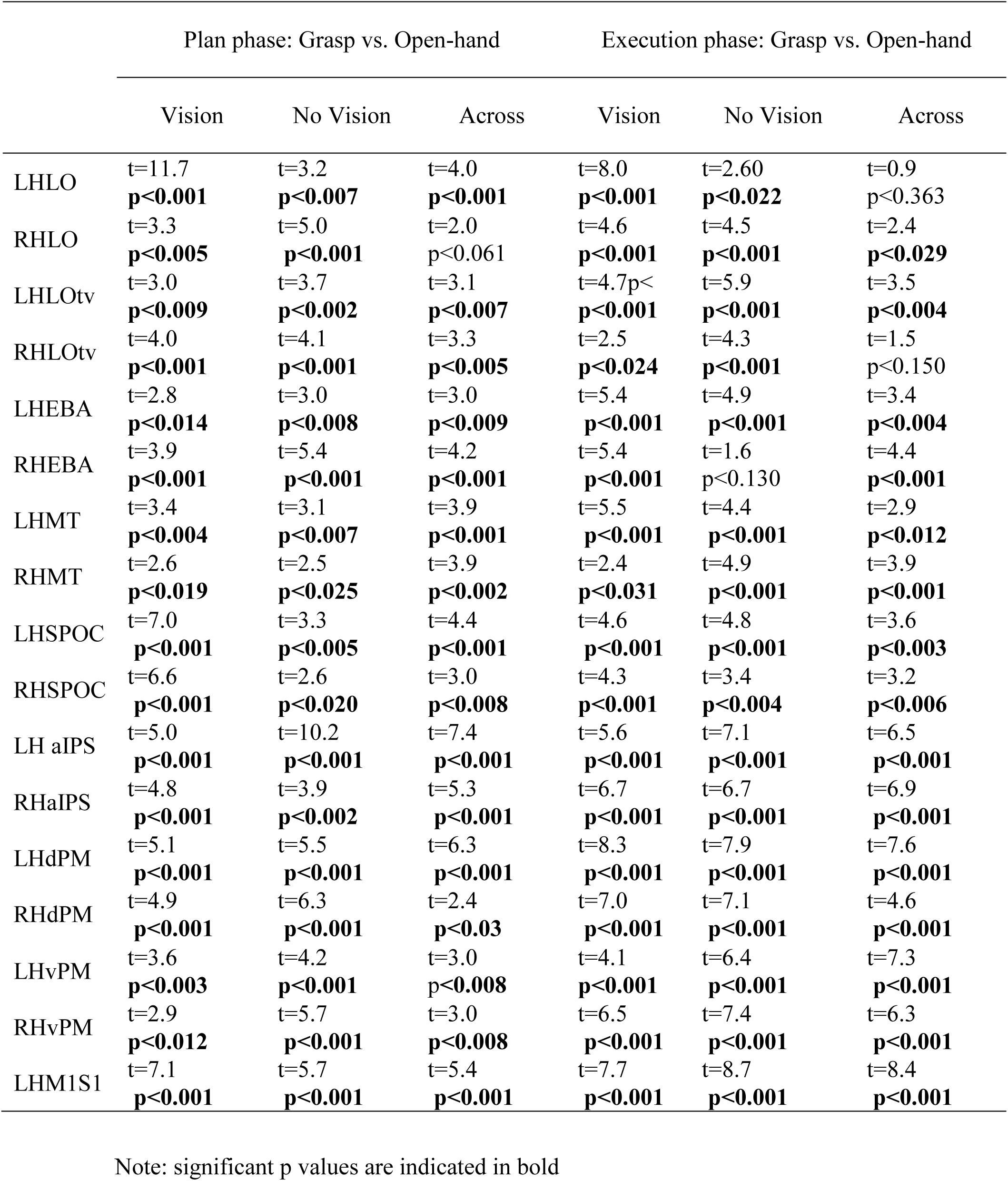
Statistical values for MVPA

|  | Plan phase: Grasp vs. Open-hand |  |  | Execution phase: Grasp vs. Open-hand |  |  |
| --- | --- | --- | --- | --- | --- | --- |
|  | Vision | No Vision | Across | Vision | No Vision | Across |
| LHLO | t=11.7<br>**p<0.001** | t=3.2<br>**p<0.007** | t=4.0<br>**p<0.001** | t=8.0<br>**p<0.001** | t=2.60<br>**p<0.022** | t=0.9<br>p<0.363 |
| RHLO | t=3.3<br>**p<0.005** | t=5.0<br>**p<0.001** | t=2.0<br>p<0.061 | t=4.6<br>**p<0.001** | t=4.5<br>**p<0.001** | t=2.4<br>**p<0.029** |
| LHLOtv | t=3.0<br>**p<0.009** | t=3.7<br>**p<0.002** | t=3.1<br>**p<0.007** | t=4.7p<<br>**p<0.001** | t=5.9<br>**p<0.001** | t=3.5<br>**p<0.004** |
| RHLOtv | t=4.0<br>**p<0.001** | t=4.1<br>**p<0.001** | t=3.3<br>**p<0.005** | t=2.5<br>**p<0.024** | t=4.3<br>**p<0.001** | t=1.5<br>p<0.150 |
| LHEBA | t=2.8<br>**p<0.014** | t=3.0<br>**p<0.008** | t=3.0<br>**p<0.009** | t=5.4<br>**p<0.001** | t=4.9<br>**p<0.001** | t=3.4<br>**p<0.004** |
| RHEBA | t=3.9<br>**p<0.001** | t=5.4<br>**p<0.001** | t=4.2<br>**p<0.001** | t=5.4<br>**p<0.001** | t=1.6<br>p<0.130 | t=4.4<br>**p<0.001** |
| LHMT | t=3.4<br>**p<0.004** | t=3.1<br>**p<0.007** | t=3.9<br>**p<0.001** | t=5.5<br>**p<0.001** | t=4.4<br>**p<0.001** | t=2.9<br>**p<0.012** |
| RHMT | t=2.6<br>**p<0.019** | t=2.5<br>**p<0.025** | t=3.9<br>**p<0.002** | t=2.4<br>**p<0.031** | t=4.9<br>**p<0.001** | t=3.9<br>**p<0.001** |
| LHSPOC | t=7.0<br>**p<0.001** | t=3.3<br>**p<0.005** | t=4.4<br>**p<0.001** | t=4.6<br>**p<0.001** | t=4.8<br>**p<0.001** | t=3.6<br>**p<0.003** |
| RHSPOC | t=6.6<br>**p<0.001** | t=2.6<br>**p<0.020** | t=3.0<br>**p<0.008** | t=4.3<br>**p<0.001** | t=3.4<br>**p<0.004** | t=3.2<br>**p<0.006** |
| LH aIPS | t=5.0<br>**p<0.001** | t=10.2<br>**p<0.001** | t=7.4<br>**p<0.001** | t=5.6<br>**p<0.001** | t=7.1<br>**p<0.001** | t=6.5<br>**p<0.001** |
| RHaIPS | t=4.8<br>**p<0.001** | t=3.9<br>**p<0.002** | t=5.3<br>**p<0.001** | t=6.7<br>**p<0.001** | t=6.7<br>**p<0.001** | t=6.9<br>**p<0.001** |
| LHdPM | t=5.1<br>**p<0.001** | t=5.5<br>**p<0.001** | t=6.3<br>**p<0.001** | t=8.3<br>**p<0.001** | t=7.9<br>**p<0.001** | t=7.6<br>**p<0.001** |
| RHdPM | t=4.9<br>**p<0.001** | t=6.3<br>**p<0.001** | t=2.4<br>**p<0.03** | t=7.0<br>**p<0.001** | t=7.1<br>**p<0.001** | t=4.6<br>**p<0.001** |
| LHvPM | t=3.6<br>**p<0.003** | t=4.2<br>**p<0.001** | t=3.0<br>**p<0.008** | t=4.1<br>**p<0.001** | t=6.4<br>**p<0.001** | t=7.3<br>**p<0.001** |
| RHvPM | t=2.9<br>**p<0.012** | t=5.7<br>**p<0.001** | t=3.0<br>**p<0.008** | t=6.5<br>**p<0.001** | t=7.4<br>**p<0.001** | t=6.3<br>**p<0.001** |
| LHM1S1 | t=7.1<br>**p<0.001** | t=5.7<br>**p<0.001** | t=5.4<br>**p<0.001** | t=7.7<br>**p<0.001** | t=8.7<br>**p<0.001** | t=8.4<br>**p<0.001** |

Note: significant p values are indicated in bold

#### Plan phase

We found significant decoding of action type both within (yellow and red bars) and across (orange bars) visual conditions in LOtv, EBA, MT, SPOC, aIPS, dPM, and vPM bilaterally, as well as in LO and M1/S1 in the left hemisphere. Area LO in the right hemisphere showed significant decoding accuracy in Vision and No Vision condition but not across visual conditions. In addition, there was higher decoding accuracy in Vision than No Vision conditions in left LO, higher decoding accuracy in Vision than cross-decoding in left LO, M1\S1 and bilateral SPOC, and higher decoding accuracy in No Vision than cross-decoding in right LO, dPM and vPM.

#### Execution phase

We found significant decoding of action type both within (blue and light green bars) and across (dark green bars) visual conditions in bilateral LOtv, MT, SPOC, aIPS, dPM, vPM and M1/S1, as well as in right LO and left EBA. Right LOtv showed above chance decoding accuracy for the dissociation between Grasp and Open hand movements in Vision and No Vision conditions but not across visual conditions. Right EBA showed above chance decoding accuracy in Vision and across visual conditions, but not in No Vision. In addition, LO in the left hemipshere showed higher decoding accuracy in Vision as compared to No Vision and across visual conditions, and LOtv and EBA in the left hemisphere showed higher decoding accuracy in Vision and No Vision conditions as compared to across visual conditions. Area MT in the left hemisphere showed higher decoding accuracy in Vision than across visual conditions. Area LOtv in the right hemisphere, as well as dPM and vPM in the left hemisphere showed higher decoding accuracy for No Vision than across visual conditions. It is important to note that differences in brain activity patterns between Action Type (Grasp vs. Open hand) and Vision conditions (Vision vs. No Vision) during the Execution phase might reflect behavioural differences. Indeed, the behavioural analyses have shown that RTs were shorter for Grasp than Open hand movements, and in Vision as compared to No Vision conditions. As such, we focused the discussion of our results on the Plan rather than the Execution phase.

#### Control areas

We ran the same analyses on two non-cortical control areas inside and outside the brain, that are not expected to show significant decoding between actions or visual conditions. As expected, the classification did not reveal any above chance decoding accuracy in these areas for any phase of the trial (Supplemental Figure 1).

## Discussion

Our results show two main findings. First, during the planning phase action intention can be reliably decoded in areas of dorsal and ventral visual stream regardless of the availability of visual information, as well as across visual feedback conditions. This suggests that in addition to vision, also proprioceptive, somatosensory, and memory-based information play a role in movement planning in dorsal as well as ventral stream areas. Second, during the execution phase we could decode actions in dorsal and ventral stream areas even when no vision was available. Further, we could dissociate action types across visual conditions, suggesting that the activity pattern was modulated by the motor output and online somatosensory feedback.

### Information driving preparatory signals in dorsal and ventral visual stream areas

In line with previous findings, our results show that the activity pattern in ventral stream areas during the planning phase preceding the action is modulated by the upcoming movement when visual feedback is available and the object is identical for both action types (Gallivan 2013, Freud 2108). This suggests that the critical factor that induces this effect is related to the visual anticipations of the upcoming action rather than the object itself. Indeed, the two movements elicit different hand configurations that lead to different perceptual expectations. In line with this, neurophysiology studies have recently shown that pre-movement neuronal activity in dorsal stream area V6A, of which SPOC is likely the homolog (Cavina-Pratesi et al. 2010a; Monaco et al. 2011; Pitzalis et al. 2013), is influence by vision of an object when a subsequent movement towards the object is required, but not in passive viewing tasks (Breveglieri et al. 2016; Fattori et al. 2017). Moreover several studies have shown that the neuronal activity in macaque’s intraparietal area is influenced by vision of the object before as well as during a grasping task (Taira et al. 1990; Sakata et al. 1995; Murata et al. 1996, 2000). An alternative and non-exclusive possibility is that the ventral stream contributes with dorsal stream areas to integrate the visual goal into the action plan, and therefore the object is processed in an action-dependent manner. In fact, grasping requires detailed information about object size and shape, while open hand movements do not.

Interestingly, we found that the activity pattern in ventral stream areas is also modulated by the upcoming action during the planning phase in No Vision conditions, and therefore driven by memory-based processes even in absence of any visual feedback. Ventral stream areas are known to have a crucial role in delayed actions that rely on memory-based information about the object, and consistently show reactivation during delayed actions performed in the dark (Fiehler et al. 2011; Singhal et al. 2013; Monaco et al. 2017). Further, patients with visual agnosia caused by lesions in the ventral stream show impairments in delayed grasping performed towards objects that are no longer visible (Milner and Goodale 1995), and transcranial magnetic stimulation in LOC impairs delayed but not immediate actions (Cohen et al. 2009). This evidence indicates a causal role of ventral stream areas in delayed actions, and our results extend previous findings by showing that the memory-based representation in ventral stream areas is not limited to the execution of the action, but encompasses also the planning phase that precedes the action. This memory-based information likely consists of the expected and well-known visual, somatosensory and motor responses generated by an action, as well as the proprioceptive inputs. Indeed, we are aware of this information even if we close our eyes.

Importantly, we could reliably decode action types in most ventral stream areas during the preparatory phase also across visual conditions. Crucially, the common aspect between the two visual conditions was action preparation. In fact, the same actions were being prepared during the delay regardless of the presence of visual guidance. One possible explanation for these results is that cross-decoding reflects the integration of proprioceptive inputs about the current state of the hand into a forward model for the upcoming action towards the target. This seems plausible since proprioceptive inputs and final target locations remained unchanged in visual and no visual conditions. Although proprioceptive information per-se might not be processed in the ventral stream, it might contribute to building a perceptual and multisensory representation of limb position (Mercier et al. 2008). This explanation is supported by evidence showing that ventral stream area EBA has a causal role in combining perceptual information about body posture and desired action goal (Zimmermann et al. 2016) for the computation of forward models and state estimation that are part of action planning (Wolpert and Ghahramani 2000). Additional evidence showing functional connections between area LO and S1 during passive tactile stimulation of the hand in absence of any movement or motor task (Tal et al. 2016), might support an alternative explanation related to the anticipation of somatosensory consequences of the action via connections with dorsal stream areas. The second and non-exclusive possibility is that the cross-decoding reflects motor imagery related to the action, which might be present in vision and no vision conditions. Indeed, the neural processing of visual imagery is evident regardless of whether participants have their eyes open (Cichy et al. 2012) or closed (Kosslyn and Thompson 1995). Also, visual imagery can induce motion aftereffects regardless of whether participants have their eyes open or closed (Winawer et al. 2010). However, since our task involved the preparation of an action, the imagery component, if present, was likely more motor than visual.

Visual and motor imagery might engage different neural mechanisms. Indeed, unlike visual imagery, the cortical representation of motor imagery, especially in the occipito-temporal cortex, has given mixed results (Oosterhof et al. 2012). For instance, proprioceptive information plays an important role in motor imagery. In fact, it has been shown that peripheral proprioceptive information about hand posture induces stronger motor facilitations during motor imagery tasks when the hand posture is compatible with the imagined movement as opposed to when it is not (Vargas et al. 2004; Fourkas et al. 2006). Strong evidence comes from deafferented subjects who do not show any motor facilitation during motor imagery because of the lack of afferent feedback (Mercier et al. 2008). Despite the possibility that motor imagery might have driven our cross-decoding results, it is unlikely that participants were engaging in motor imagery for simple actions, such as grasping an object. Indeed, motor imagery requires a conscious effort, while planning a simple hand movement is a process that we perform effortlessly. In addition, planning a real movement and imagining one are processes that differ at several levels, as the former involves real consequences while the latter does not. Therefore, our cross-decoding possibly tackles information that is specific to anticipations of real movements in both visual conditions rather than motor imagery.

The decoding accuracy in Vision condition during the planning phase was higher than cross-decoding in LO, SPOC and interestingly in dPM and M1/S1. Additionally, LO showed higher decoding accuracy in Vision than No Vision condition. These results indicate a strong visual component in areas that are adjacent to the occipital lobe, such as LO and SPOC, that might be implicated in processing the visual anticipations of movements at the early stage of the planning phase. Subsequently, this information might be sent forward to premotor and motor areas for their integration in the motor plan at a later stage. The lack of a sensory specialization for the anticipation of actions in other ventral and dorsal stream areas is likely due to the fact that anticipations of the consequences of an action are based on experience, and during the execution of actions in our everyday life, visual and somatosensory responses do not happen in isolation. In fact, they are naturally synchronized so that every visual frame of our hand movement corresponds to a specific somatosensory feedback. Thus, at the neural level, the strong experience-based association of visual and somatosensory feedback that we build throughout life might reinforce functional connections between ventral and dorsal stream areas. Indeed, evidence from neurophysiology and neuroimaging studies have shown anatomical and functional connections between dorsal and ventral stream areas (Borra et al. 2008; Verhagen et al. 2008; Bracci et al. 2012; Hutchison and Gallivan 2018), as well as between the primary somatosensory cortex and area LO in the ventral stream (Tal et al. 2016). These connections might allow cross-talks not only during movement execution, but also during the planning phase that precedes an action.

### Feedback generated during action execution

Neurophysiology works have shown that dorsal stream areas are specialized in processing visual and somatosensory information for action, as well as proprioceptive information about arm location (Sakata et al. 1973; Lacquaniti et al. 1995). Therefore, it is not surprising that we can reliably use the activity pattern in areas of the dorsal stream to decode actions based on visual, somatosensory and proprioceptive feedback during action execution. Indeed, the known multisensory aspect of the parietal and frontal cortices makes of the dorsal stream a valuable site to integrate information coming from different senses (Buneo and Andersen 2006). Since experience allows for accurate anticipations, it might be the case that this information is also used in these areas to anticipate the consequences of upcoming actions during the planning phase. Although the pattern classifications during the execution phase might reflect behavioural differences in reaction times between actions types, the finding of a strong memory-based and somato-motor representation of actions performed with no visual guidance in ventral and dorsal premotor cortex is in line with neurophysiology studies showing that grasping actions elicit robust neuronal activity in macaque ventral and dorsal premotor cortex regardless of whether the action is performed with or without visual guidance (Raos et al. 2004, 2006).

Actions, and in particular grasping movements, also require a perceptual component in order to identify the target object that we intend to interact with as well as the hand approaching the object. Given the known role of human ventral stream areas in perceptual identification of shapes (Kourtzi and Kanwisher 2000, 2001; James et al. 2003), body parts (Downing et al. 2001), and hands (Bracci et al. 2010), it is expected that visual information about any of these categories would modulate the activity pattern in ventral stream areas during the execution of the movement when vision is available. Therefore, the action-driven modulation of the activity pattern during action execution in ventral stream areas can easily be explained by the difference in visual feedback elicited by Grasp vs. Open hand movements. In addition, behavioural differences in RTs between action types might also be reflected in differential brain activity patterns.

One of the surprising findings of this study is that even very simple actions, like grasping repeatedly the same object or open the hand on top of it, engage neural mechanisms that can be used to distinguish between two ordinary movements. These results offer evidence that planning actions is a pervasive process that involves many brain areas, regardless of how effortless the action might be. Complex actions, novel contexts and dynamic situations likely require efforts that might drive higher decoding accuracies as compared to simple movements. Yet, natural actions are part of our everyday repertoire. It is enough to think about how smoothly we grasp our cup of coffee on the desk, reach to press the elevator button, or open our hand above the eyes to shade from the sun. Despite of how fluent and automatic actions are, many brain regions reflect the preparation needed to perform the movement.

## Conclusions

Predictions are at the basis of skilled behaviors as well as simple hand actions. Although we perform grasping movements effortlessly, the motor plan is supported by a myriad of information that spans the visual, somatosensory, motor and cognitive systems. Therefore, the prediction of upcoming movements is based on several types of neural signals that we normally experience on a daily basis. Despite the specialization of ventral stream areas in visual recognition of shapes and dorsal stream areas in somatosensory and motor processing, we used the activity pattern in both ventral and dorsal stream areas to reliably decode action preparation based on brain signals evoked by visual and non-visual (kinesthetic and motor) information. Our results show that visual, memory and somato-motor information similarly affect the pattern of activity in areas of dorsal and ventral visual streams for the representation of the motor plan, and allows predicting action intention across domains, regardless of the availability of visual information. In summary, the strong associations between visual and somato-motor information in our everyday life allows predicting action intention based on visual and non-visual inputs in dorsal and ventral visual streams.

## Ethical Approval

All procedures performed in this study were in accordance with the ethical standards of the Human Research Ethics Committee of the University of Trento (protocol 2016-021) and with the 1964 Helsinki declaration and its later amendments or comparable ethical standards.

## Acknowledgements

The authors would like to thank Pietro Chiesa for technical support, Jason Gallivan for providing the EBA and LO localizers and Angelika Lingnau for providing the MT localizer.

