## Supplemental Material for "Predictive coding of action intentions in dorsal and ventral visual stream is based on visual anticipations, memory-based information and motor preparation"

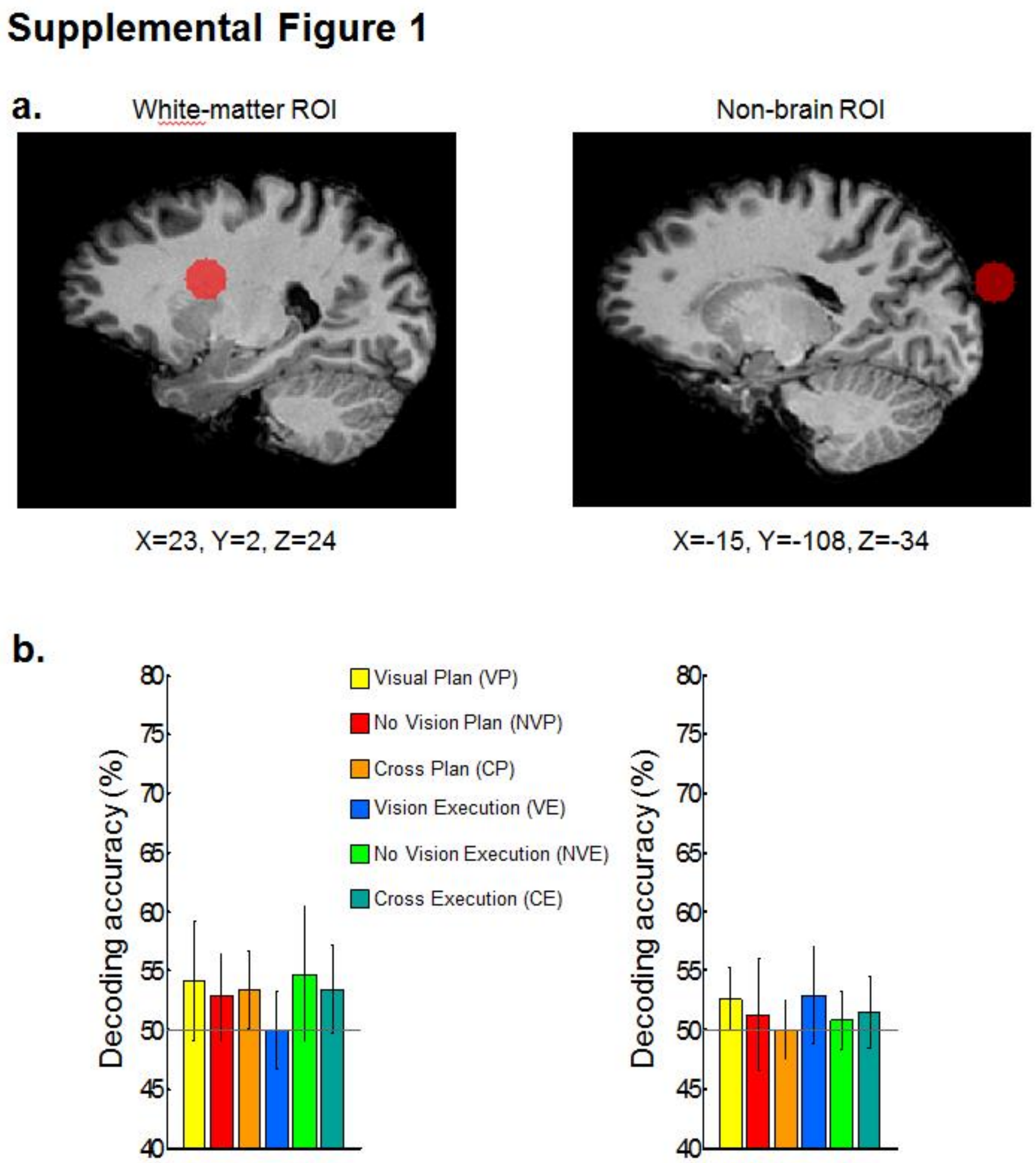


***Supplemental Figure 1***

Classifier decoding accuracies in non-brain control regions. a) Non-brain control

ROIs. b) Classifier accuracies for Plan and Execution phases for the right white-matter ROI (left) and non-brain ROI (right). Chance level is indicated with a line at 50% of decoding accuracy. Error bars show 95% confidence intervals. Importantly, no significant differences were found with respect to 50% chance.

### Supplemental Figure 2

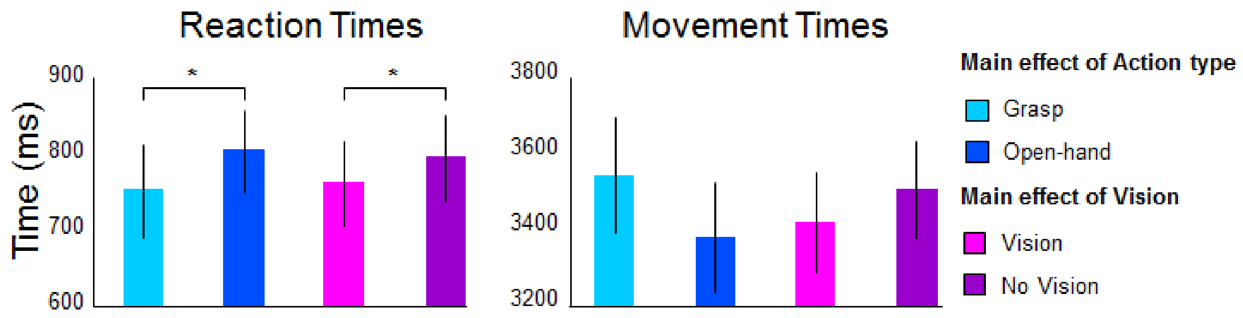


***Supplemental Figure 2***

Bargraphs of the Reaction Times (left panel) and Movement Times (right panel). The outcome of the ANOVA is indicated by the asterisk: *p<0.05. Only significant results are indicated. The error bars represent the 95% confidence intervals.

***Supplemental Table 1. Average reaction and movement times***

| | Grasp | Open hand | Vision | No Vision |
|---|---|---|---|---|
| RTs | 752 | 805 | 762 | 796 |
| MTs | 3547 | 3381 | 3420 | 3507 |

**Statistical methods for Univariate analyses**

First, we explored the extent to which the preparatory responses in ventral and dorsal stream areas are shaped by the upcoming action when the object is visible (in Vision condition) as well as when the object is not visible and has to be retrieved from memory (in No Vision conditions). To this aim, we performed our analysis on the Plan phase preceding the movement. Second, we examined which brain areas are more strongly influenced by the view of the hand approaching the object (in Vision condition) as compared to performing actions in the dark (in No Vision conditions). To this goal, we performed the analysis in the Execution phase. Notably, the involvement of brain areas in action planning and execution in Vision and No Vision conditions indicated by univariate analysis, include a somatosensory component elicited by predictive mechanisms, in the planning phase, and the movement itself, in the execution phase.

For each ROI we used SPSS to perform one ANOVA for each phase of the task: Action Plan and Execution. Each ANOVA had two levels for action type (Grasp and Open-hand) and two levels for availability of vision (Vision and No Vision). When interactions reached significance, we further explored the nature of the interaction by performing post-hoc two-tailed t-tests for four comparisons: Grasp Vision vs. Open Hand Vision, Grasp No Vision vs. Open Hand No Vision, Grasp Vision vs. Grasp No Vision, Open Hand Vision vs. Open Hand No Vision.

To control for multiple comparisons, a false discovery rate (FDR) correction of $q \leq 0.05$ was applied, based on the number of ROIs and number of t-tests performed within each time phase (Benjamini and Yekutieli 2001).

### Results of Univariate Analyses

The Talairach coordinates for each ROI are specified in Table 1. Statistical values of the main effects and post-hoc t-tests for each area are reported in Supplemental Table 2 and 3.

*Plan phase*

As shown in Supplemental Figure 3, planning grasping movements elicited higher activation than planning open hand movements during the Plan phase in left SPOC, right aIPS, vPM and LO. In addition, vision induced higher activation as compared to when no vision was available in bilateral SPOC. Conversely, bilateral EBA, left vPM and M1/S1 showed higher activation for No Vision than Vision conditions. Specifically, the ANOVA on the activation levels during the Plan phase showed a main effect of action type in left SPOC, right aIPS, vPM and LO with higher activation for Grasp than Open hand movements. Further, there was a main effect of Vision in left SPOC, bilateral EBA and left M1/S1, with higher activation for Vision than No Vision in left SPOC and the opposite pattern in bilateral EBA, left vPM and M1/S1.

*Action phase*

As shown in Supplemental Figure 3, the execution of a grasping movement elicited higher activation than open hand movement in bilateral aIPS and dPM, left M1/S1, SPOC, EBA, LOtv, and MT. In addition, all areas, except left M1/S1 and bilateral vPM, showed higher activation in Vision than No Vision conditions. Specifically, we found a main effect of action type in bilateral aIPS and dPM, left M1/S1, SPOC, EBA, LOtv, and MT, as well as a main effect of vision in bilateral aIPS, dPM, SPOC, EBA, LO, LOtv, and MT. In addition, there was a significant interaction in left aIPS, LO, MT, vPM and right dPM. However, post-hoc comparisons survived

FDR correction ($p<0.011$) only for left aIPS, vPM and right dPM. In particular, the interaction in left aIPS was due to higher activation for Vision than No Vision conditions during the Open hand but not the Grasp movement. Given the well-known role of this area in grasping movements, it is likely that the execution of a grasp fully engaged area aIPS regardless of the presence of vision, while the activation for Open hand movements was enhanced when the hand was visible as compared to when it was not. In addition, the interaction in left vPM and right dPM was related to higher activation for Grasp than Open hand in No Vision but not in Vision conditions.

### Supplemental Figure 3

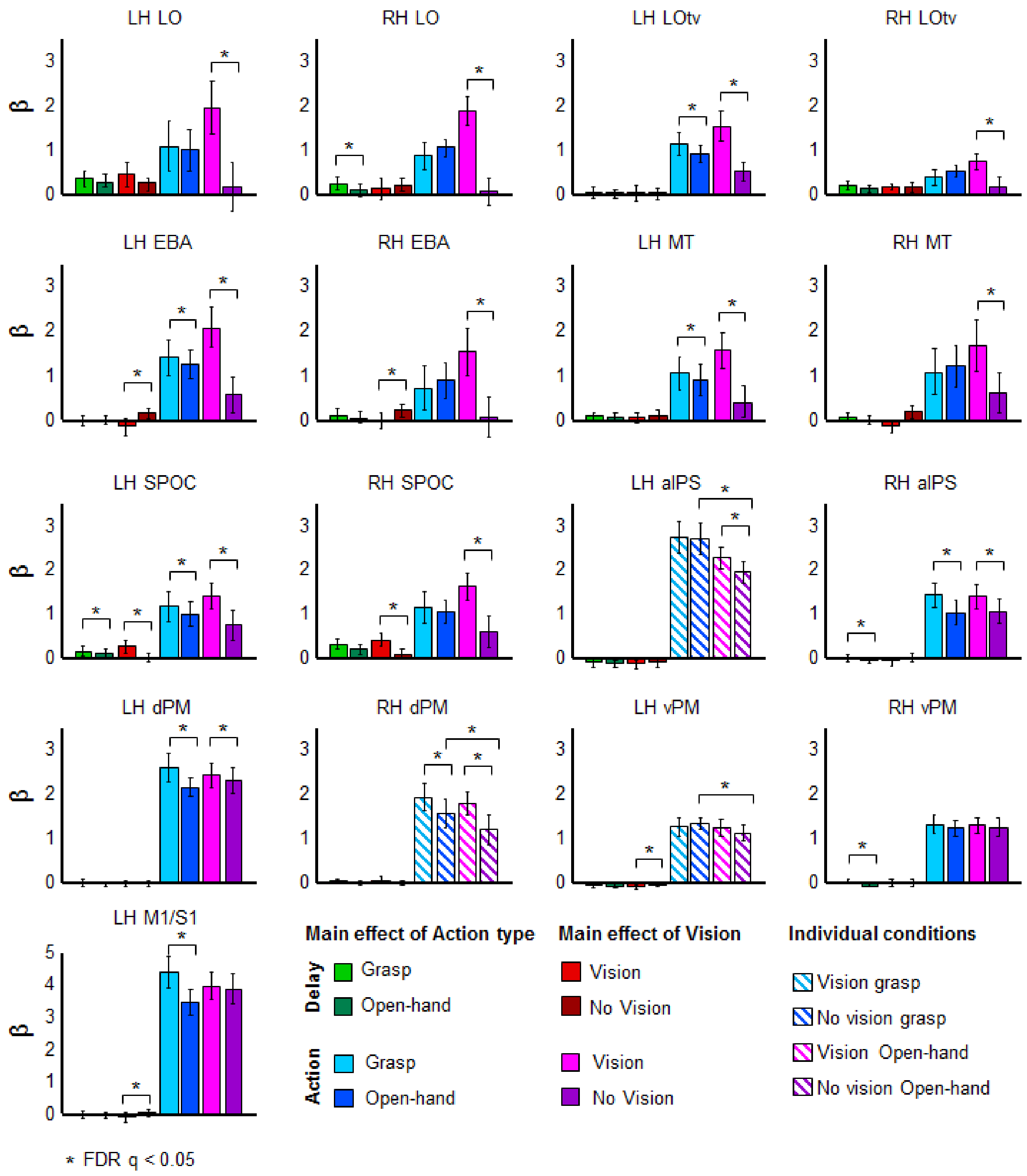

***Supplemental Figure 3***

Activation levels in our ROIs. For each area, the bar graph indicate the β weights for the main effects of Action type (Grasp and Open-hand) and Vision (Vision and No Vision) in the planning phase (left) and execution phase (right). When interactions reached significance, post-hoc t-tests are shown between β weights of individual experimental conditions. Error bars show 95% confidence intervals.

***Supplemental Table 2. Statistical values for main effects and interactions in univariate results***

| | Plan phase | | | | Execution phase | | | | | |
|---|---|---|---|---|---|---|---|---|---|---|
| | ME action type | | ME vision | | ME action type | | ME vision | | Action type by vision | |
| | $F_{(1, 15)}$ | p | $F_{(1, 15)}$ | p | $F_{(1, 15)}$ | p | $F_{(1, 15)}$ | p | $F_{(1, 15)}$ | p |
| LHLO | | | | | | | 65.4 | 0.001 | 19.6 | 0.001 |
| RHLO | 5.2 | 0.038 | | | | | 83.7 | 0.001 | | |
| LHLOtv | | | | | 12.6 | 0.003 | 29.1 | 0.001 | | |
| RHLOtv | | | | | | | 24.1 | 0.001 | | |
| LHEBA | | | 7.2 | 0.017 | 4.9 | 0.04 | 42.2 | 0.001 | | |
| RHEBA | | | 5.2 | 0.037 | | | 64.2 | 0.001 | | |
| LHMT | | | | | 8.6 | 0.01 | 66.4 | 0.001 | 8.5 | 0.01 |
| RHMT | | | | | | | 28.7 | 0.001 | | |
| LHSPOC | 4.7 | 0.046 | 11.4 | 0.004 | 7 | 0.018 | 29.3 | 0.001 | | |
| RHSPOC | | | | | | | 52.9 | 0.001 | | |
| LH aIPS | | | | | 33.4 | 0.001 | 9.5 | 0.008 | 11.6 | 0.004 |
| RHaIPS | 4.6 | 0.049 | | | 8.6 | 0.01 | 66.4 | 0.001 | | |
| LHdPM | | | | | 18.7 | 0.001 | 2 | 0.172 | | |
| RHdPM | | | | | 16.6 | 0.001 | 29.6 | 0.001 | 4.6 | 0.049 |
| LHvPM | | | 5.339 | 0.035 | | | | | 6.7 | 0.02 |
| RHvPM | 10.2 | 0.006 | | | | | | | 6.3 | 0.02 |
| LHM1S1 | | | 8.8 | 0.01 | 83.5 | 0.001 | | | | |

Note: Only significant results are shown

***Supplemental Table 3. Statistical paired-sample t-tests for univariate results***

| | Grasp > Open hand | | Vision > No Vision | |
|---|---|---|---|---|
| | Vision | No Vision | Grasp | Open hand |
| LH aIPS | | | | t=4.1 p<0.001 |
| RHdPM | | t=3.8 p<0.002 | | |
| LHvPM | | t=2.9 p<0.011 | | |

Note: Only significant results are shown
